# Malathion-resistant *Tribolium castaneum* has enhanced response to oxidative stress, immunity, and fitness

**DOI:** 10.1101/2021.11.16.468861

**Authors:** Abdur Rauf, Richard M Wilkins

## Abstract

Many cases of insecticide resistance in insect pests give resulting no-cost strains that retain the resistance genes even in the absence of the toxic stressor. Malathion has been widely used against the red flour beetle, *Tribolium castaneum* Herbst. in stored products although no longer used. Malathion specific resistance in this pest has provided resistance that is long lasting and widely distributed. To understand this resistance a malathion resistant strain was challenged with a range of stressors including starvation, hyperoxia, malathion and a pathogen and the antioxidant responses and some lifecycle parameters were determined.

Adult life span of malathion-specific resistant strain of *T. castaneum* was significantly shorter than the susceptible. Starvation and/or high oxygen reduced adult life span of both strains. Starving with and without 100% oxygen gave longer lifespan for the resistant strain, but for oxygen alone there was no difference. Under oxygen the proportional survival of the resistant strain to the adult stage was significantly higher, for both larvae and pupae, than the susceptible. The resistant strain when stressed with malathion and/or oxygen significantly increased catalase activity, but the susceptible did not. The resistant strain stressed with *Paranosema whitei* infection had significantly higher survival compared to the susceptible, and with almost no mortality.

The malathion resistant strain of *T. castaneum* showed greater vigour than the susceptible in most oxidative stress situations and especially where stressors were combined. The induction of the antioxidant enzyme catalase could have helped the resistant strain to withstand oxidative stresses, including insecticidal and importantly those from pathogens. These adaptations, in the absence of insecticide, seem to support the increased immunity of host insects to pathogens seen in other insect species, such as mosquitoes. By increasing the responses to a range of stressors the resistant strain could be considered as having enhanced fitness.

## 1. Introduction

Malathion resistance in the red flour beetle, *Tribolium castaneum* (Herbst), is a worldwide problem and as it is very stable it became widespread in natural populations (Dyte and Blackman, 1970), with the first case of malathion-specific resistance in *T. castaneum* reported in Nigeria (Parkin et al. 1962). In the absence of insecticide, this resistance does not reduce fitness, being the genetic contribution to future generations. Evidence for malathion specific resistance shows, that in the absence of treatment, there is no fitness difference between resistant and susceptible strains or the resistant strain has a fitness advantage (Beeman and Nanis 1986; McKenzie and Farrell 1993; White and Bell 1995; Haubruge and Arnaud 2001; Arnaud et al. 2002; Arnaud and Haubruge 2002; Bughio and Wilkins 2004; Arnaud et al. 2005; Okoye et al. 2007; Kliot and Ghannin 2012; Grigoraki et al, 2017). A malathion-specific resistant strain showed an 8-23% increase in biotic potential (fecundity and developmental time) relative to the susceptible strain (Haubruge and Arnaud 2001). Malathion-specific carboxyesterases (MCE) have been implicated in a range of insect species. These include several strains of mosquitoes (Whyard and Walker, 1994; Whyard et al. 1995), moths (Beeman and Schmidt 1982), pteromalid parasitoids (Baker et al. 1998), beetles (Price 1984, Spencer et al. 1998, Haubruge et al. 2002) and fruit flies (Wang et al, 2017). The persistence of the malathion resistance in these species/strains is often not known.

However, most studies have shown fitness costs associated with insecticide resistance (McKenzie and Farrell 1993; Kliot and Ghannin 2012; Cao et al., 2014; Guillem-Amat et al., 2020), and specifically for *T. castaneum* in the case of phosphine resistance (Oppert et al., 2015; Nayak et al., 2020). In fact, in 60% of fitness reports there was a cost for resistance but less commonly for organochlorines (Freeman et al, 2021).

Exposure to radiation, ultraviolet light, and some chemicals results in the production of reactive oxygen species (ROS) such as hydrogen peroxide and superoxide anion radical, usually via the mitochondria of cells (Towarnicki et al., 2020). In general, ROS are harmful to living organisms because ROS can cause oxidative damage to proteins, nucleic acids and lipids (Gladyshev 2014; Sies, 2015). In this context, ROS has been recognized to be related to animal aging and life span (Parkes et al., 1999; Otali, 2014; Zhang et al., 2017). ROS stimulates signal transduction (Schieber and Chandel, 2014) and mediates various responses such as cell growth and apoptosis (Suzuki et al. 1997; Speakman et al, 2015). On the other hand, ROS plays a helpful role in the innate immunity system of an insect (Futo et al, 2016; Khan et al, 2017). Hydrogen peroxide may have a role in effecting hormesis (Ludovico and Burhans, 2014). Living organisms thus require a regulatory system for ROS. Antioxidant enzymes scavenge ROS. Catalase (H_2_O_2_ oxidoreductase; EC 1.11.1.6, CAT) plays an essential role and catalyses the degradation of H_2_O_2_ to water and oxygen (Yamamoto et al. 2005; Sies, 2017).

Other antioxidant enzymes such as superoxide dismutase, glutathione transferase, and glutathione reductase have been characterized in insects (Felton and Summers, 1995). Oxidative stress results from an imbalance of oxidants and antioxidants, either a surfeit of oxidants and/or a deficit of antioxidants (Gladyshev 2014). The antioxidant enzymes superoxide dismutase (SOD), catalase (CAT), and glutathione peroxidase (GPx) are the main enzymatic defences and act in concert with a panoply of non-enzymatic antioxidants (Kodrík et al, 2015). The elaboration of antioxidant defences is presumed to have played a critical role in the evolution of oxygen-respiring organisms, especially terrestrial arthropods. Catalase, by virtue of its ability to break down H_2_O_2_ into H_2_O and O_2_, plays an important role in mitigating oxidative damage, since H_2_O_2_ is a ready source for hydroxyl radical (.OH) formation via Fenton chemistry. Together with superoxide dismutase (SOD), catalase forms a major antioxidative axis, and, as such, can impact directly on cellular redox status (Klichko et al., 2004; Sies, 2017).

GSTs play a vital role in protecting tissues against oxidative damage and oxidative stress. Elevated GSTs in a resistant strain attenuated the pyrethroid-induced lipid peroxidation and reduced mortality, whereas their inhibition eliminated their protective role (Vontas et al., 2001).

Stressors which increase ROS in insects can include starvation, elevated oxygen levels, toxins (e.g. malathion) and parasites/pathogens. We will consider the effects of these on a susceptible and a malathion-resistant strain of *T. castaneum*. Various classes of pesticides induce reactive oxygen species and oxidative tissue damage which may contribute to the toxicity of these xenobiotics (Abdollahi et al., 2004). Reactive oxygen species may serve as common mediators of programmed cell death (apoptosis) in response to many toxicants and pathological conditions (Bagchi et al., 1995). In the case of *Drosophila melanogaster*, treatment with malathion increased stress tolerance and life span (i.e. fitness) (Bonilla et al., 2002). Low doses of imidacloprid bind to the insect brain, causing rapid release of Ca^+^ and ROS (Martelli et al., 2020). The insect is weakened and less able to tolerate environmental stresses. In fact, catalase in this species decreases with age and plays a role in redox signalling (Lennicke and Cochemé, 2020).

Oxidative damage in insects is thought to be high because they usually have no oxygen binding proteins in their blood so oxygen is carried directly to their cells via the tracheolar network which exposes them to four-to-five-fold higher concentrations of oxygen than in animals with blood haemoglobin. Free radicals impair metabolism and necessitate protection and repair especially during starvation, poor diet, desiccation and rehydration. Raising houseflies in an atmosphere of 100% oxygen reduced their mean and maximum life span and it increased their levels of protein carbonyls in whole body extracts (Sohal et al., 1993) and in the mitochondria (Towarnicki et al., 2020). Cells exposed to low oxygen levels (hypoxia) activate the transcription factor hypoxia-inducible factor 1 (HIF-1) as an adaptive response. Cells exposed to hypoxia do not undergo senescence or cell death and do not diminish ATP levels. By contrast, cells exposed to high oxygen levels (hyperoxia) undergo senescence and cell death and decrease their ATP levels, yet do not activate HIF-1 (Harrison et al., 2006). Despite these divergent responses with respect to senescence, cell death, metabolism, and gene expression, the signalling events in both systems are mediated by the generation of mitochondrial-derived reactive oxygen species (ROS) (Chandel and Budinger, 2007). High mortality of *T. castaneum* was found when exposed to 90% oxygen (Calderon, 1991; Wang et al., 2018). Adult *T. castaneum* under hypoxia depressed mitochondrial function but following reoxygenation increased their antioxidant activity (Wang et al., 2018). Long term selection of *Drosophila melanogaster* with 90% oxygen generated a resistant strain that were larger and had increased body weight by 20% (Zhao et al., 2010).

In view of the increased fitness of the highly malathion-specific resistant strain of *T. castaneum* characterised here, and the universal replacement of previous susceptible populations by similar malathion resistant strains (Attia et al., 2020), it was relevant to determine its responses to additional oxidative stresses. Previous work had shown enhanced fitness when this strain was stressed on a less suitable grain (rice) for food (Bughio and Wilkins, 2004). Further studies showed enhanced gut capability in terms of tolerance to protease inhibition (Bughio and Wilkins, 2021). The role of catalase, the crucial antioxidant enzyme (Sies, 2017), was to be explored, using the highly resistant strain. Although malathion is little used for *T. castaneum* management, the characteristics of resistant strains, which are widespread in stored products (Anusree et al, 2019), are important in developing sustainable strategies. Understanding the redox relationship in the resistant strain is useful when considering the use of phosphine, other insecticides/natural products or natural enemies in any extant population of this pest (Nayak et al., 2019). Thus, the objectives were to compare the responses of an insecticide resistant strain, with no-cost fitness traits and which is widespread, to stressors such as hyperoxia, starvation, pathogen and malathion, and combinations (with a standard susceptible strain). Responses included longevity, survival, immunity and catalase induction, and their contributions to fitness.

The choice of parasite stressor was *Paranosema whitei* (Weiser) (species previously known as *Nosema whitei* Weiser has now been replaced (Sokolova et al., 2005)). This is an important parasite for *Tribolium* and a potential biocontrol agent, and is an obligate, intra-cellular microsporidian (protozoan). The pathogen has been almost always lethal in the late larval and pupal stages of the host *T. castaneum* so that only a few individuals manage to enclose to adult (Milner, 2006). Transmission of *P. whitei* is horizontal and occurs after host death when the infected larva, pupa or adult dies, and is destroyed either when the host medium is mechanically treated (as in stored products) or the carcass is cannibalized.

## 2. Materials and methods

### 2.1 Insect strains *red flour beetle T. castaneum*

Two strains of *T. castaneum*, PH-1 and FSS-II, were maintained at the School of NES, Newcastle University, UK. The malathion-specific resistant Ph-1 strain (Rauf and Wilkins, 2002; Haubruge et al, 2002) was previously received from Natural Resources Institute, UK and the FSS-II strain of *T. castaneum* was obtained from Central Science Laboratory, York, UK. The susceptible strain of *T. castaneum*, FSS-II, was described by Lagisz et al. (2010). Wholemeal wheat flour supplemented with 5 % brewer’s yeast at 29±1° C and 65±5 % RH, in the dark, was used as the culture medium. The natural resistant strain was further selected with malathion by topical application of technical grade malathion to the adults to give around 90% mortality and the survivors were used to produce the next generation. This treatment was repeated for every third generation. After five such sequential selections, the initial LD_50_ value of 6.4 μg/beetle increased to 135 μg/beetle (Ph-5) and after 10 selection cycles the LD_50_ had risen to 162 μg/beetle (Ph-10). The enhanced strains were utilized in conjunction with the reference susceptible FSS-II strain. These field derived strains were used to ensure stable and effective resistance levels and although they are not directly genetically-related they provide realistic subjects. Responses to sequential malathion selection are given in Table 1. Further characterisation of these strains with respect to other insecticides and synergists are provided in Supplementary Materials.

**Table 1.** LD_50_ values (and 95% confidence intervals) and resistance factors (RF) of susceptible and resistant strains of *T. castaneum*

| Strain<sup>a</sup> | LD<sub>50</sub> µg/insect | Slope ±SE | RF |
| --- | --- | --- | --- |
| FSS-II | 0.027 (0.02-0.03) | 3.92±0.51 | - |
| Ph-1 | 6.4 (5-8) | 2.76±0.32 | 237 |
| Ph-2 | 75 (65-94) | 4.81±0.61 | 1562 |
| Ph-5 | 135 (112-170) | 4.84±0.55 | 2812 |
| Ph-10 | 162 (142-191) | 5.15±0.57 | 3375 |

<sup>a</sup> Ph strain, number of malathion selection cycles (every 3 generations) starting with Ph-1.

### 2.2 Application of oxidative stress to *T. castaneum* using oxygen

#### 2.2.1 Adult lifespan

Pure oxygen atmosphere conditions were maintained in a 30×30×30 cm clear plastic chamber, kept within an incubator, by passing a continuous gentle stream of humidified filtered 100 % oxygen (from a gas cylinder BOC) through the chamber with the outlet bubbled through water outside the chamber. A 29±1° C temperature and 65±5 % relative humidity was maintained in the chamber. To study the effect of pure oxygen on the adult stage of the strains of *T. castaneum*, 2-5-day-old beetles (20) of each strain were kept in 30 ml glass vials (5 in total) with culture medium (1 g) in each, within the chamber. The food medium was changed every 15 days. In the control, filtered humidified normal air (supplied from a compressed air cylinder BOC) was used instead of pure oxygen. The effect on the adult mortality was observed with and without providing food to the beetles. Under fed conditions, mortality was noted every two days under oxygen and every fortnight in the air/control; while, where the beetles were kept starved, mortality was observed every day in both oxygen stressed and control conditions. The data obtained were analysed using survival curves drawn using Graphpad Prism V9 to obtain number of days for median survival of the population and curve comparison under the various stress conditions.

#### 2.2.2 Short-term stress to determine catalase activity

Adults of both strains were stressed with malathion; 100 adult beetles were treated topically (prior to placing in the chamber) with a technical grade malathion solution in acetone at LD50 level (adults placed in 1g culture medium, 3 replicates). Mortality was noted after 24 h with, and without, pure oxygen. The survivors were killed by freezing and kept at -20° C to determine catalase activity.

#### 2.2.3 Survival of immature stages

The survival of larval and pupal stages of the strains of *T. castaneum* was noted by counting the number of adults that emerged after 15 days in the case of larvae and 10 days for the pupae. Ten last instar larvae or newly pupated pupae were transferred to vials containing culture medium (1 g) and kept in the chambers with air or 100% oxygen under standard conditions. For each life stage five replicates were used.

### 2.3 Determination of catalase activity

#### 2.3.1 Homogenisation of insects

Crude enzyme extracts of *T. castaneum* were prepared by homogenising 40 mg of adult beetles in 1.5 ml of ice cold 0.05 M phosphate buffer pH 7. The homogenates were centrifuged and the supernatants were used as the enzyme source.

#### 2.3.2 Assay of catalase activity

Catalase activity was determined by using a modification of the spectrophotometric method of Durusoy et al. (1995). Reactions, at 30° C, were initiated by the addition of 50 µl of the supernatant to 3 ml of a substrate solution containing 0.05 M sodium-potassium phosphate buffer, pH 7, and H2O2 at a final concentration of 20 mM. Catalase activity was measured as a decrease in H2O2 concentration. The decrease in absorbance at 230 nm was monitored for 1.5 min and was noted at every 0.5 min. The consumption rate of H2O2 was calculated using a calibration curve of H2O2 in a concentration range from 2 to 50 mM. The initial activity was determined from the slope of the line (Δ O.D. per min). The activity was expressed as micromole of H2O2 decomposed per min per milligram of protein. Protein concentration of the homogenates was determined by the method of Bradford (1976) using bovine serum albumin as a standard.

### 2.4 Paran*osema whitei* infection of *T. castaneum*

The malathion-specific resistant Ph-5 and susceptible FSS-II strains of *T. castaneum* were used to test their susceptibility to *Paranosema whitei*, a microsporidian pathogen of this species. An infected stock of FSS-II strain was the inoculum (Khan and Selman, 1988). To prepare the infective material containing the pathogen spores, the host larvae killed by the disease were ground with a mortar and pestle. The dried crushed material was sieved through 100-mesh sieve and the powder was stored at room temperature to use for infection.

For each strain, ten 30 ml glass vials were prepared, each containing wholemeal wheat flour (10 g) with 5% brewer’s yeast. Twenty randomly selected same-age beetles of each strain, irrespective of gender, were put in each vial for egg laying and kept at 29±1° C and 65±5 % relative humidity. After three days the beetles were removed by sieving. The above infective powder (30 mg), containing disease spores, was added and carefully mixed in each vial. The emergence of adult beetles was noted after 35 days.

### 2.5 Statistical analysis

The response data were analysed by probit/logit regression analysis using the computer programme POLO PC (Leora Software, Berkeley, CA, 1987). All other results were analysed with GraphPad Prism V9 for Windows (GraphPad Software San Diego, CA, USA) using two-way analysis of variance ANOVA, with posttest Bonferroni correction. Survival curves were drawn using Graphpad Prism V9 using the Kaplan-Meier survival fractions. Survival curves were compared using the Log-rank (Mantel-Cox) test with pre-planned pairs.

#### 3. Results

### 3.1 Exposure of adult insects to oxidative stress

#### 3.1.1 Adult life span of the strains of *T. castaneum* with, and without, pure oxygen conditions

The results show that adult life-spans of both strains were significantly reduced (*P*<0.0001) under high oxygen conditions, as compared to the control in normal air (Table 2), irrespective of other stressors. However, starved insects were less impacted by oxygen exposure (medians reduced to 0.40 and 0.45 of control for the susceptible and resistant strains, respectively) than fed insects (median values reduced to 0.22 and 0.28 of control) with the resistant strain relatively less affected in both nutritional conditions (Table 2).

**Table 2.**
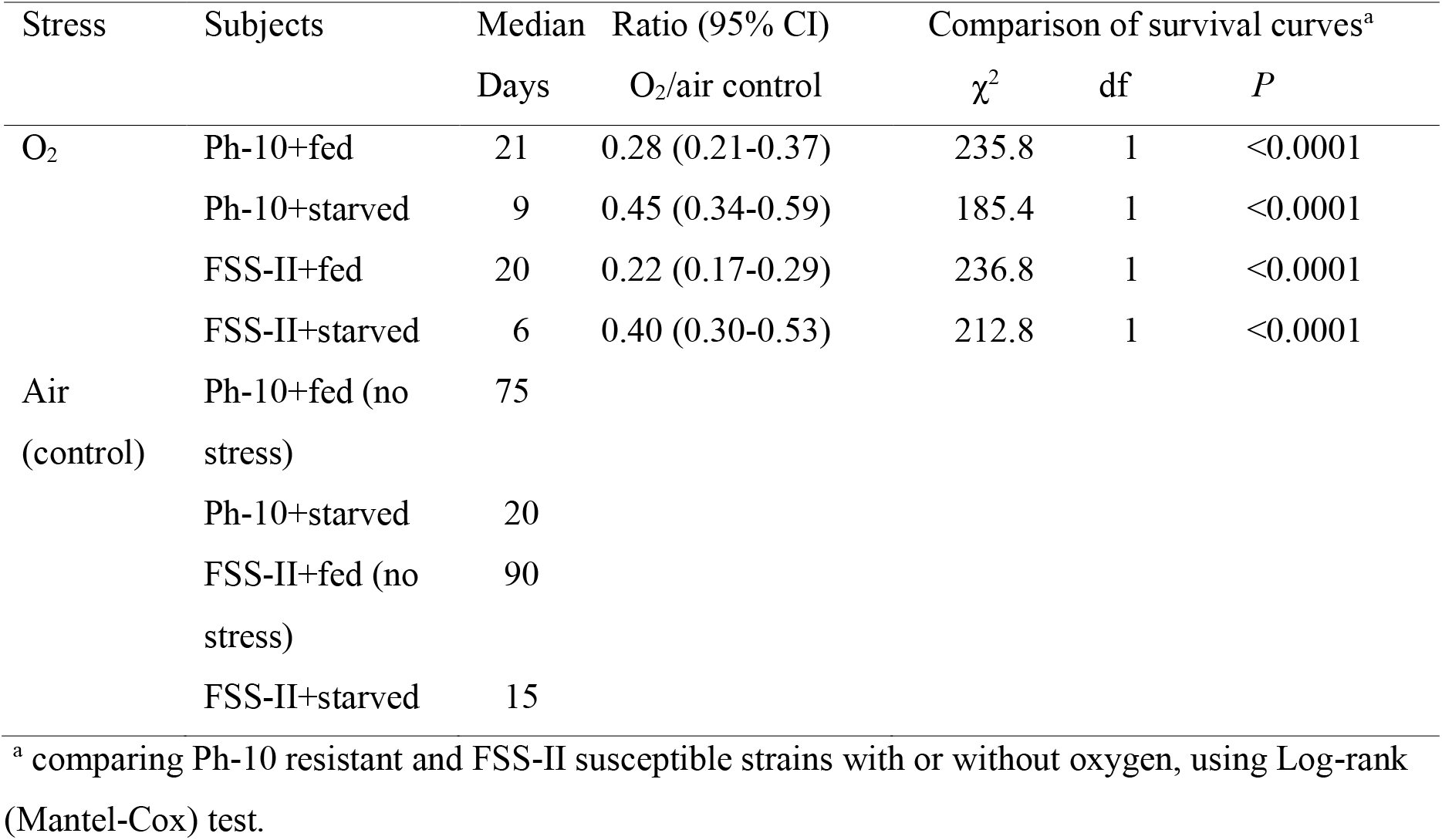
Survival of the adults of strains of *T. castaneum*, comparing the responses to pure oxygen

In comparing the two strains, under fed conditions, the susceptible FSS-II strain survived significantly better (*P* <0.0001) under normal air conditions as compared to the malathion-specific resistant strain of *T. castaneum* Ph-10 (Fig 1 and Table 3). Under 100% oxygen this was reversed (*P*<0.05), with the resistant strain living slightly longer (Fig 1 and Table 3) although the difference between the two was significant.

**Fig. 1.**
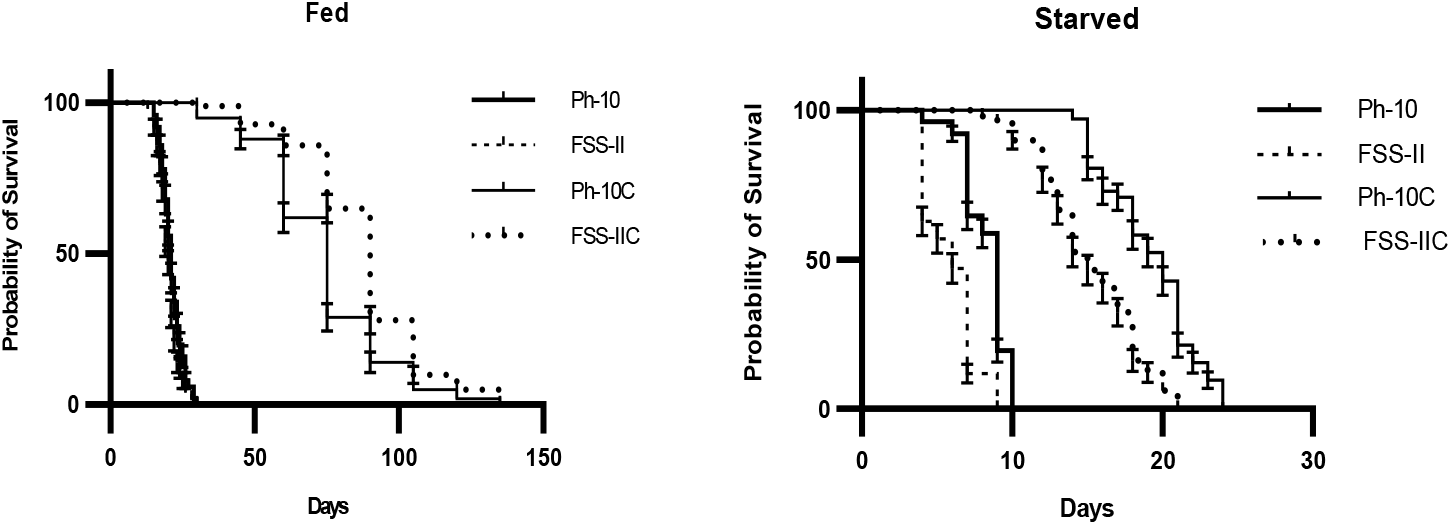
Survival in adults of the strains of *T. castaneum* where wheat flour was used as food (Fed), and where no food was used (Starved) with 100% oxygen (Ph-10 and FSS-II) or control with air (Ph-10C and FSS-IIC). Ph-10 resistant, FSS-II susceptible. Error bars represent SEM.

**Table 3.** Summary of survival of the adults of strains of *T. castaneum* and comparison of survival curves

| Strain | Stress | Median | Ratio R/S<br>and (95% CI) | Comparison of survival curves<sup>a</sup> |  |  |
| --- | --- | --- | --- | --- | --- | --- |
| | | Days | | $\chi^2$ | df | *P* |
| Ph-10 | air+fed (no stress) | 75 | 0.83 (0.63-1.10) | 17.83 | 1 | <0.0001 |
|  | air+starved | 9 | 0.60 (0.46-0.79) | 185.4 | 1 | <0.0001 |
|  | oxygen+fed | 21 | 1.05 (0.80-1.38) | 4.96 | 1 | 0.026 * |
|  | oxygen+starved | 9 | 1.50 (1.14-1.98) | 75.94 | 1 | <0.0001 |
| FSS-II | air+fed (no stress) | 90 |  |  |  |  |
|  | air+starved | 15 |  |  |  |  |
|  | oxygen+fed | 20 |  |  |  |  |
|  | oxygen+starved | 6 |  |  |  |  |

<sup>a</sup> comparing Ph-10 resistant (R) with FSS-II susceptible (S), using Log-rank (Mantel-Cox) test.
\* significantly different at <5% *P*

The starved individuals of both strains lived significantly shorter (P<0.0001) lives than fed insects in both 100% oxygen and the normal air control environment (Fig 1 and 2). Under starved conditions in normal air, the susceptible strain had shorter adult lives than the resistant. However, if pure oxygen is added as an extra stressor, the median survival for both is lowered substantially, but the resistant strain is less damaged.

**Fig 2.**
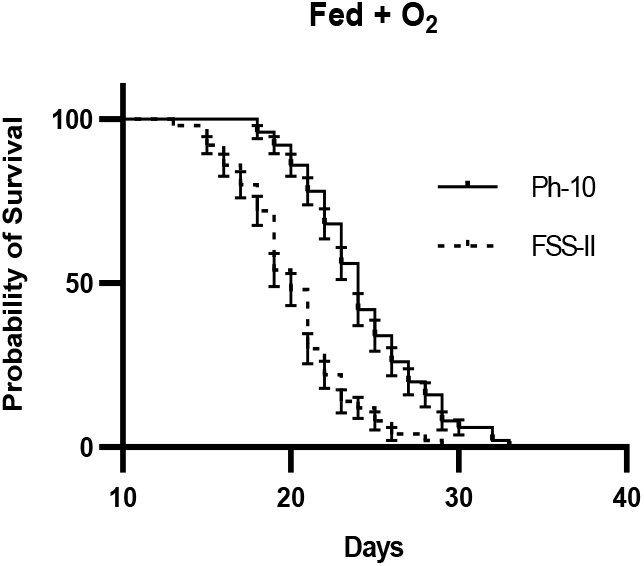
Survival in adults of the strains of *T. castaneum* where wheat flour was used as food with 100% oxygen. Ph-10 resistant, FSS-II susceptible. Error bars represent SEM. Ph-10 curve displaced +2.

#### 3.1.2 Survival of the late larval & pupal stages to adulthood of the strains of *T. castaneum* maintained under air and pure oxygen conditions

Both the larval and pupal stages of the susceptible strain of *T. castaneum* survived slightly better than the resistant in normal air conditions, but this difference was statistically non-significant (Fig 3). Comparing the two strains for late larval and pupal stages using Bonferroni correction gave, for air (control) conditions, larvae; t (32)=0.40, P > 0.05 (*NS*), pupae; t (32)=1.12, *P* > 0.05 (*NS*). Under oxygen stress conditions, the survival percent of the resistant Ph-10 strain to the adult stage was significantly higher, both in larvae (t (32)=4.45, *P* <0.001), and pupae (t (32)=2.42, *P* < 0.05), than the susceptible FSS-II strain (Fig 3). This enhanced survival of the no-cost malathion-resistant strain over the susceptible is consistent through the main stages of the life cycle.

**Fig. 3.**
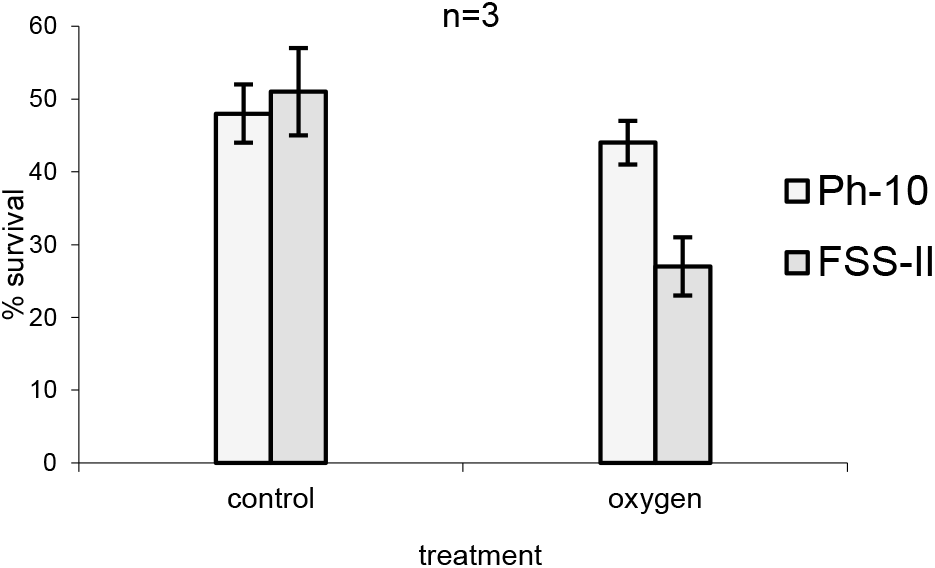
The survival (%) in late larval and pupal stages, to adult, in air (control) or 100 % oxygen for the strains of *T. castaneum*. n= 5, number of replications, where 10 individuals for each replication were used. Ph-10 resistant, FSS-II susceptible.

#### 3.1.3 Adult mortality in the two strains of *T. castaneum* treated with malathion in air or pure oxygen

The mortality in the resistant Ph-10 strain adults after topical treatment with malathion at LD_50_ level was marginally higher (t (8)=0.49, non-significant P>0.05, Bonferroni post-test) than the susceptible FSS-II strain where treated beetles were kept in normal air (Fig. 4). This slight variation can be explained by dosing errors. However, when combining the stressors (insecticide plus oxygen), the malathion-treated resistant strain showed significantly better survival (t (8)=2.83, P<0.05) when kept under the pure oxygen conditions (24 h) after treatment with malathion at the same LD_50_ level.

**Fig. 4.**
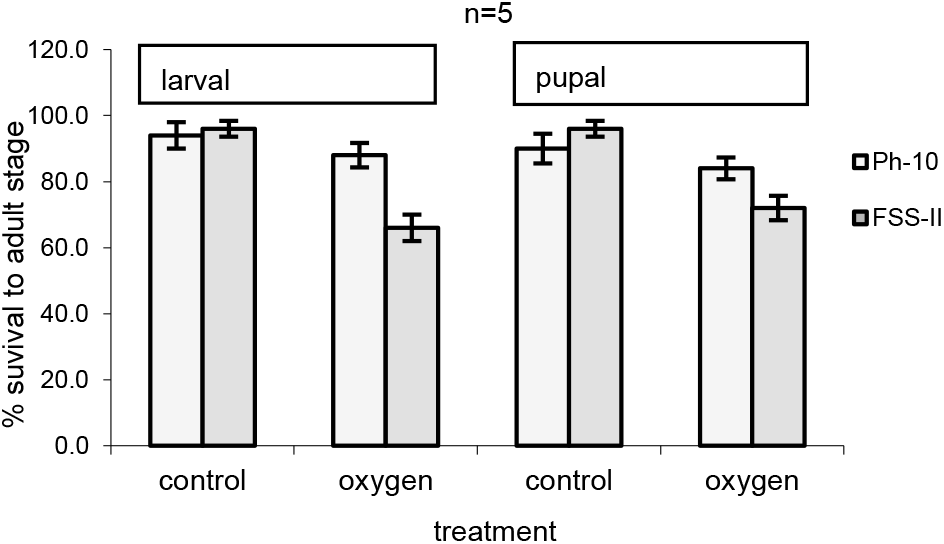
Survival of adults of the strains of *T. castaneum* after treatment with malathion at LD_50_ level and without, and with, 24 hours 100% oxygen. n=3 number of replications, where 100 beetles for each replication were used.

#### 3.1.4 Catalase activity in the strains of *T. castaneum* after malathion and pure oxygen treatment

The catalase activities in non-insecticide-treated individuals of both strains were not significantly different from control following oxygen exposure (Fig. 5) (resistant Ph-10 t (16)= 1.77, P>0.05) and susceptible FSS-II (t (16)= 0.55, P>0.05). The mean catalase activity was not increased, in the malathion resistant strain after topical treatment with malathion (t (16)= 0.92, P>0.05, NS), as compared to the untreated, where the beetles were kept under normal air conditions. For the susceptible strain there was also no significant change (t (16)= 0.22, P>0.05). But, the increase, in the enzyme activity, was significantly higher, than the control, for the resistant strain (t (16)= 2.54, P<0.05), in the malathion treatment with oxygen for 24 hours. There was no significant change in the enzyme activities observed in the malathion susceptible strain (t (16)= 0.14, P>0.05), both after separate and combined effects of malathion and oxidative stress. Catalase activity was determined 24 h after acute treatment with malathion.

**Fig. 5.**
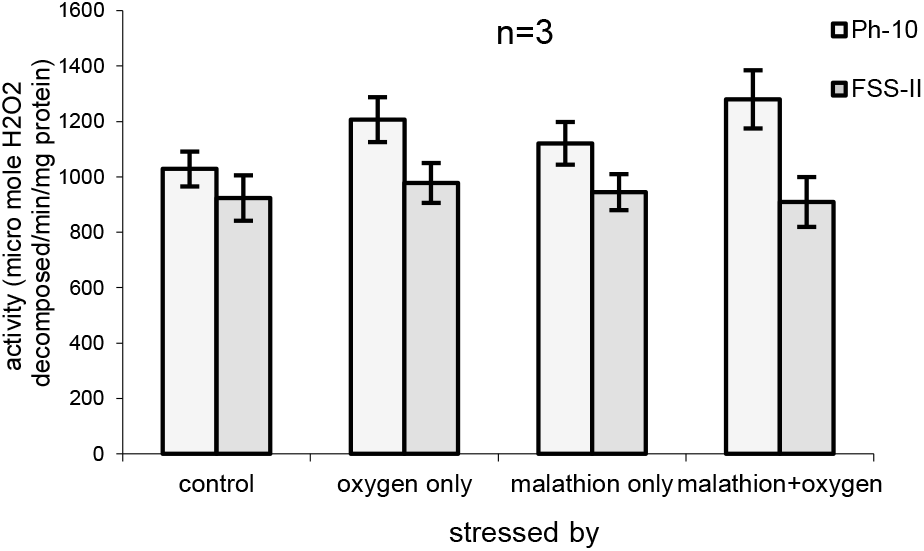
Catalase activity in the strains of *T. castaneum* after stress with malathion at LD_50_ level and 24 hours 100% oxygen. n=3 number of replications, where, 40 mg of beetles for each replication were used.

#### 3.1.5 Paranosema whitei infection of T. castaneum

Following infection of the early larval stage of the strains of *T. castaneum* with *Paranosema whitei*, we compared those beetles that grew to adulthood. The adult survival rate in the resistant (Ph-5) strain was very significantly greater (t (36)=10.48, P<0.001) than in the susceptible (FSS-II) strain (t (36)=0.27, P>0.05) (Figure 6). There was little disease effect on the resistant strain (P>0.05) while the susceptible strain was nearly eliminated.

**Fig. 6.**
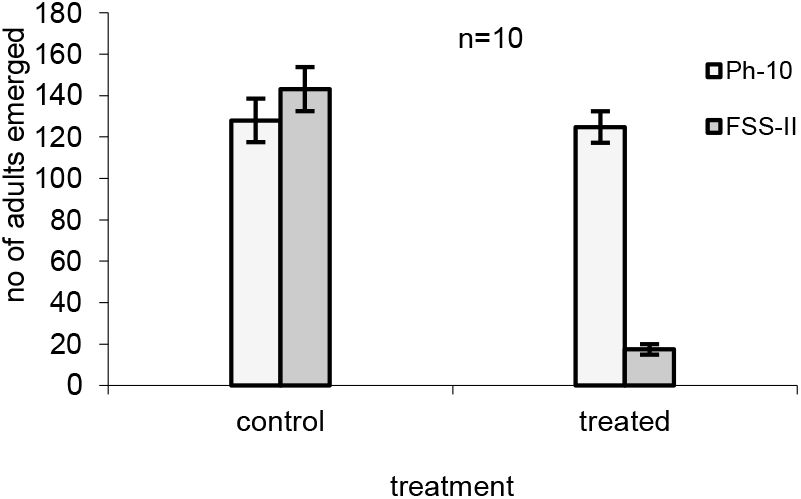
Adult survival after treatment with *Paranosema whitei* in the strains of *T. castaneum*. n=10 number of replications, where, 20 beetles for each replication were used. Errors bar represent SEM.

## 4. Discussion

### 4.1 Lifespan and survival of the strains exposed to oxidative stressors

Lifespan or longevity in adults of *T. castaneum* under laboratory conditions can extend from 60 to 200 days when strains developed under varying temperatures were compared (Hani Soliman and Lints, 1982). Depending on the strain, larval growth rates were negatively correlated with longevity of the adults. Strains originating from grain stores were compared for adult longevity and mean survival times were found to vary from 120 to 170 days (Kavallieratos et al, 2020) when fed on cracked barley. When fed on unsuitable rice grain the longevity was drastically shortened. All these insects were reared on wheat flour previously to the study (Kavallieratos et al, 2020). Generally, the expected adult lifespan under controlled conditions does depend on the strain (and conditions); however, there are no published results for insecticide-resistant strains of this species, particularly a no-cost resistant strain. The maximum lifespans of the strains in our study, which also had been in culture for many years, were 135 days (both strains) in fed conditions, in line with the previous studies. Under starved conditions, lifespans fell to 24 days (resistant Ph-10) and 21 days (susceptible FSS-II). These reductions are similar to non-resistant adults when fed on the unsuitable rice diet (Kavallieratos et al, 2020). Adult malathion-resistant *Musca domestica* when exposed to oxygen survived better than a susceptible strain and induced higher catalase levels (Rauf et al, 2008).

In our study we found that the adults of highly malathion-specific resistant strain of *T. castaneum* had relatively shorter lives (based on median values), in the normal air environment, than the susceptible and thus the resistant strain was less fit on the basis of adult life span. Presumably this could make the resistant strain adults less damaging in a store, in suitable low stress conditions. However, the diverse stress conditions (moisture, temperature, quinone levels, pathogens, etc) in a grain store environment needs to be considered. This shorter lifespan may not translate to decreased reproduction over generations but the susceptible strain has been replaced universally in grain stores by the malathion resistant strain in spite of this (Beeman and Nanis, 1986; Arnaud et al, 2005; Attia et al, 2020). Previous research on similar *T. castaneum* strains showed egg hatch percentages and larval weights were significantly lower in the malathion-resistant strain compared to the susceptible strain but on less suitable rice flour the malathion-resistant strain laid more eggs than the susceptible (Bughio and Wilkins, 2004). Fitness benefits of the resistance in this strain become apparent when stressed, which stress is not insecticidal, but not in an unstressed environment.

In a comparison of phosphine susceptible and resistant *T. castaneum* genes associated with the mitochondria had elevated expression in resistant insects (Oppert et al., 2015) but phosphine resistance is usually not stable (Djihinto et al., 2013). Similar shorter lifespans have been found for an artificially enhanced permethrin resistant strain of *Anopheles gambiae*, which had a knockdown resistance (kdr) mutation and enhanced levels of P450 and esterase enzyme activities. This shows that lifespan effects are not limited by the nature of the insecticide, including resistance mechanisms (Otali et al, 2014).

The malathion-resistant strains of *T. castaneum* have exhibited fitness advantages in the absence of insecticide (Haubruge and Arnaud, 2001, Arnaud and Haubruge, 2002, Bughio and Wilkins, 2004, Arnaud et al, 2005), whereas, the majority of fitness trials (which usually exclude adult lifespan) have shown that there are fitness costs associated with other insecticide resistant species. The proportion of fitness cost resistance is 60% of published research (Freeman et al, 2020). Malathion-resistant strains of stored-product insects often show no reduced fitness compared to susceptible in the absence of any selection pressure (White and Bell 1995, Mason 1998, Guedes et al, 2017).

As mentioned previously that some malathion-resistant *T. castaneum* strains expressed fitness advantages in the absence of malathion (ffrench-Constant and Bass, 2017; Steinbach et al, 2017). However, life span had not been assessed as a component of these fitness studies and the enhanced fitness in the malathion specific strain was due to improved fecundity and reproductive potential (Arnaud et al., 2002). This is an important area as the mechanisms for fitness costs provide the basics of economic optimal models of insecticide resistance management (Brown et al., 2013).

Oxygen, as a highly oxidizing molecule, generates free radicals mainly via the mitochondria (Otali et al., 2014), which then participate in other oxidative chemical reactions (Finkel and Holbrook, 2000; Abele, 2002; Harrison et al., 2006) that are directly correlated with degenerative processes, illnesses and mortality (Skulachev, 2002; Cheng et al., 2003). We have noticed a significant reduction in adult lifespan, in an oxygen atmosphere, of both malathion resistant and susceptible strains of *T. castaneum* which confirms the above findings (Calderon et al, 1991) as well as the relationship with age (Lee and Ducoff, 1983). Sohal et al (1993) reported that raising houseflies in an atmosphere of 100% oxygen reduced their mean and maximum life span and it increased their levels of protein carbonyls in whole body extracts. Generating 90% oxygen resistant *Drosophila melanogaster* by selection over 13 generations lead to gene upregulation and increased body weight (Zhao et al., 2010). A similar response in evolved increased body size was also seen by Klok et al (2009). These fly strains were not assessed for any insecticide resistance. We also have found that the resistant strain Ph-10 was better able to endure oxidative stress and lived significantly longer, in 100% oxygen environment, than the susceptible FSS-II. Moreover, during oxidative stress conditions, the percent survival, of both late larval and pupal stages to adulthood of the malathion-specific resistant strain, was higher than the susceptible strain. Further oxidative stress was caused by starving the insects and this was related to a reduction in the lifespans, again less in the resistant strain. Fitness was reduced by both stressors and this has been confirmed by transcriptome analysis in *T. castaneum* (Koch and Guillaume, 2020). A related result has been shown with 1 % lyophilized broccoli, included in the food, which significantly elongated the life span of the (not resistant) red flour beetle (*T. castaneum*) under physiological conditions (32 °C) and under heat stress (42 °C) (Grunwald et al, 2013). Similarly, *T. castaneum* adults when stressed by heat had extended survival with grape seed extract in their diet (Grunwald et al, 2014). Surprisingly, food grade antioxidants were harmful to *T. castaneum* when mixed with food; however, the diet was peanut kernels (Garcia et al, 2017).

### 4.2 The endogenous antioxidant enzyme, catalase, and its responses to oxidative stressors in the strains of *T. castaneum*

It is known that most insecticides, including malathion, produce oxidative stress in tissues through the formation of reactive species (ROS) (Abdollahi et al., 2004; Possamai et al., 2007; Guedes et al, 2016; Martelli et al, 2020). The basis of organophosphate toxicity in the production of oxidative stress may be due either to (a) their “redox-cycling” activity, where they readily accept an electron to form free radicals and then transfer them to oxygen to generate superoxide anions and hence hydrogen peroxide through dismutation reactions, or (b) to ROS generation via changes in normal antioxidant homeostasis resulting in antioxidant depletion, if the requirement of continuous antioxidants is not maintained (Kovacic 2003; Yonar 2013).

The unstressed catalase levels in the two strains did not differ significantly. This is in line with Awan et al, (2012) who found no differences between deltamethrin-resistant and susceptible strains of *T. castaneum*. However, in the mosquito *Anopheles funestus*, resistant strains (to permethrin) had significantly higher catalase activities than susceptibles (Oliver and Brooke, 2016). When treated with the catalase synergist 3-amino-1, 2, 4-triazole (ATZ) there was significant increase in mortality using deltamethrin (Oliver and Brooke, 2016).

In our study we tested the effect of pure oxygen on the mortality of malathion-specific resistant and susceptible strains of *T. castaneum* treated with malathion, and subsequent *in vitro* changes in the key antioxidant enzyme catalase. We found a significant increase in mortality of the susceptible strain as compared to the resistant, under the 100% oxygen conditions which suggests that there might be an additional mechanism in the resistant strain to combat the oxidative stress, along with the confirmed carboxylesterase malathion-specific resistance mechanism (Kodrik et al, 2015; Oliver and Brooke, 2016)

Although there was no significant difference in the catalase activity between the strains, in the air control, there was an induction in the enzyme activity in the resistant strain, after treatment with malathion, with oxygen treatment or after the combined effect of malathion and oxygen, while no significant effect of the treatments was noted in the susceptible strain. Based on the known weight of adult *T. castaneum* of these strains (Bughio and Wilkins, 2021) of 2.3mg (Ph-10) and 2.2mg (FSS-II) the dosages of malathion applied can be calculated as 70430 µg g^-1^ (Ph-10) and 12.27 µg g^-1^ (FSS-II) (to achieve LD_50_, assuming complete penetration of cuticle). However, these concentrations will be substantially proportionally reduced at 24 h when catalase activity was determined. Although the total esterase activity in the resistant was lower than in the susceptible strain the specific malathionase in vivo was much higher (7.74 vs. 0.32 pmole/min) and was 28 times higher *in vitro* (see supplementary material). Thus, remaining malathion could have been below the concentration needed in both strains to elicit significant increases in catalase activity, although there were non-significant increases (whether active/inactive metabolites provide stress is not known). Using *Galleria mellonella* (single susceptible strain) and sublethal injected doses of imidacloprid a dose-response induction of catalase was found (Yucela and Kayis, 2019). At the lower doses there were no significant increases in catalase activity for up to 96 h post-treatment. Imidacloprid is likely more persistent in insect tissue than malathion (based on environmental data) but both insecticides induce catalase and other anti-oxidant enzymes (Yucela and Kayis, 2019.

We found the main difference between the strains was that the resistant strain could induce catalase activity in response to stress (either from malathion or O_2_) and induction increased with both stressors together (Fig 5), whilst the susceptible strain could not, and it also had a lower baseline level.

The increase in this antioxidant enzyme, in both insect strains, was probably a response towards increased ROS generation in organophosphate toxicity. Klichko et al. (2004) found an enhanced activity of catalase, in *Drosophila* strains, after exposure to 100% oxygen. DDT and malathion resistant individuals of two *Anopheles* species showed an increased capacity for coping with oxidative stress, by increasing glutathione peroxidase and catalase activity (Oliver and Brooke, 2016). The small catalase response in our study to either malathion or oxygen alone may be a result of insufficient stressor concentration within the insect body (when assayed). The above studies used stressors in the diet, or for long periods which exerted strong antioxidant responses. Our use of precise topical insecticide treatment provided a method to understand the dose-response relationship in induction of insect defence mechanisms. Further work is needed to explore the relationship in resistant strains.

The current results contradict those of Price et al. (1982) who reported reduced catalase activity, after phosphine treatment, in the resistant and susceptible strains of *Rhyzopertha dominica* but there was significantly less inhibition in the resistant strain as compared, while all the reference resistant strains exhibited higher enzyme activity than the reference susceptible strain. Catalase activity in the red flour beetle has been positively correlated with the strain resistance level to phosphine (Gao, 2009). In the stored product pest *Trogoderma granarium* peroxidase and superoxide dismutase levels were higher in the moderately phosphine resistant strains but the activity of catalase was only non-significantly increased (Yadav et al, 2020). However, the reductant phosphine probably produces diverse effects compared to organophosphates. Additionally, phosphine inhibits the transcription of the catalase gene in *Drosophila melanogaster* (Liu et al. 2017). It is known that inhibition of oxidative defences, such as catalase, resulted in increased susceptibility to insecticides in *Anopheles arabiensis* and *An. funestus*, indicating that oxidative defence is a vital part of insecticide resistance (Oliver and Brooke, 2016). Mosquitoes, stressed oxidatively with paraquat, become more susceptible to insecticides, in this case permethrin and DDT to which defences depend on the upregulation of the P450 system (Champion and Xu, 2018). This in turn depends on availability of catalase (Champion and Xu, 2018). Longevity in adult *D. melanogaster* has been associated with overexpression of Cu-Zn superoxide dismutase and catalase levels (Orr and Sohal, 1994).

Insecticide, including malathion, poisoning in higher animals, causing free radical damage is an important direct or indirect factor in several pathological and toxicological processes (Possamai et al, 2007). In contrast to our results, Vettraino et al. (2001) reported an extended longevity and a lower level of antioxidant enzyme activity in the paraquat resistant strain of *Drosophila* suggesting a qualitative change in the enzyme system which contributed towards the resistance against paraquat in the selected strain. Similarly, catalase activity was significantly higher (62%) in susceptible *Sitophilus granarius* compared to a phosphine-resistant strain (Bolter and Willian 1990).

### 4.3 Oxidative stress through infection

Our results appear to suggest that the malathion-specific resistant strain has a distinctive ability to survive against *Paranosema whitei*, a severe obligate pathogen of this insect species. It is reported that this parasite results in almost certain death of the infected populations of *T. castaneum*. (Blaser and Schmid-Hempel, 2005).

Defence in insects against infections and parasites is mediated through ROS, which are essential components in the immune system (Goswamy and Elrazoqui, 2021). Bacteria such as Wolbachia can affect ROS levels in the host (Zug and Hammerstein, 2015). As a pathogen it can precipitate an immune response and cause oxidative stress, whereas in coevolved symbioses, the relationship promotes redox homeostasis. It is known that insecticide stress impacts various immune mechanisms (James and Xu, 2012). It has been shown that in *Drosophila* an extracellular immune-regulated catalase (IRC) mediates a key host defence system that is needed during host-microbe interaction in the gastrointestinal tract. Strikingly, adult flies with severely reduced IRC expression show high mortality rates even after simple ingestion of microbe-contaminated foods (Ha et al., 2005).

Most fitness studies on insecticide resistance and pathogen stress have used mosquitoes, as their role in vectoring human disease pathogens can be crucial. In the mosquito, *Culex pipiens*, organophosphate resistant individuals had a greater load of bacteria *Wolbachia* than the susceptibles, showing a greater fitness cost (Berticat et al, 2002). In a mosquito (*Culex quinquefasciatus*) with high level of an esterase resistance mechanism the transmission of filariasis was strongly reduced (McCarroll and Hemingway, 2002). In *Anopheles gambiae* Plasmodium reduced survival in the insecticide resistant strains but not in a susceptible and there was an increase in fecundity for all strains (Alout et al, 2016). Resistance may have changed the interaction between plasmodium and the mosquito, resulting in increased fitness cost of infection. Research on *Culex pipiens* and the obligate parasite *Vavraia culicis*, which kills its host before adult eclosion compared organophosphate resistant and susceptible strains (Agnew et al, 2004). The parasite did not affect survival of the susceptible but 40% of a resistant strain died. The mechanism of resistance in this strain was enhanced esterases (Agnew et al, 2004). This is the reverse of our results; however, the resistance mechanism here was specific malathionase (general esterases did not contribute) and this may not cause ROS production, compared to general esterases. Additionally, the resistant *T. castaneum* had high fitness whereas most of the lifecycle parameters for the resistant *C. pipiens* strain were reduced (Agnew et al, 2004). In deltamethrin-resistant *Aedes albopictus* there was a fitness cost but a decrease in its competence as a vector. However, the resistant strain still could complete the period of incubation (Deng et al, 2021). Evidence for a link between innate immunity, ROS production and insecticide (including malathion) resistance in *Aedes aegypti* involving Nrf2 signalling has been found (Bottino-Rojas et al, 2018).

A review in this area (Minetti et al, 2020) reports varied effects on Plasmodium survival according to mosquito species and resistance mechanism. These authors note that “lack of research on this topic is surprising, given the critical impact that any interactions may have on the epidemiology of malaria in an era of widespread pyrethroid resistance”. Similar sentiments could be expressed in relation to pest management in stored products where malathion resistance is prevalent. Immunity is carried forward through the beetle’s stages (Critchlow et al, 2019) but in assessing the extent of the immunity (adaptive) no account was taken of the insecticide resistance of the population used or the grain species in the store. In this case, (Critchlow et al, 2019), beetles were sampled from a store in 2013, and it is possible that malathion resistance was still extant. Results could have been influenced by this; only by comparison with a truly susceptible strain could it be assessed.

Insect responses to infection are partly mediated through ROS generated by the NOX/DUOX pathway in the gut (Kim and Lee, 2014). Immune-regulated catalase protects the gut tissue from the action of ROS (Chaitanya et al, 2016). High ROS levels resulting from overexpression of Duox can disrupt cellular function and structure; in *T. castaneum* up regulation following toxic exposure led to increased antioxidant enzymes (Gao et al, 2020). This type of redox interaction may explain the augmented levels of catalase seen in the current (unstressed) malathion-resistant strain. Immune responses in *T. castaneum* exposed to *P. whitei* (Lopez-Ezquerra et al, 2018) included gene upregulation involving non-ROS changes, but the authors considered this ROS generation a possibility.

It could be considered that the malathion-resistant strain Ph-5 is itself resistant to *P. whitei*, unlike the susceptible strain. In this case, it would have achieved immunity without having previous exposure to the pathogen, that is without immunity priming. For *T. castaneum* this has been shown in the case of infection with *Bacillus thuringiensis*, but with selecting for resistance over 11 generations (Khan et al, 2017). In the current case innate immunity rather than adaptive immunity is likely, although the distinction between the two may not be clear (Criscitiello and de Figueiredo, 2013). In formulating what the relationship between insecticide resistance and immunity to a parasite might be, it may be just the properties of the strains selected here. Further work is needed to extend the implications to other strains of *T. castaneum* and of course other species with stable rersistance, but there has been some support from mosquitoes with another parasite type (McCarroll and Hemingway, 2002). In *Myzus persicae* with multiple resistance mechanisms there was no fitness cost in reproduction or susceptibility to a pathogen (Erdos et al, 2021). Thus, the pathogen control method would be compatible with insecticides, but would not be so in our case with resistant *T. castaneum* and *P. whitei*.

GST plays a major role in detoxification and in insecticide resistance, but has been shown not to be a factor in the resistant strain Ph-10. Transcriptome analysis of a susceptible strain of *T. castaneum* comparing it with RNAi treated larvae for blocking GST activity was made (Chen et al, 2016). This also suppressed genes encoding antioxidant enzymes, such as CuZnSOD, Duox, Prx, HPX, CPO, and MCORP. The authors further suggested that these influence the gene’s function on lifespan, immunity, development and reproduction. Although this work did not use a resistant strain of *T. castaneum* it may provide some insight in how resistance to insecticides, and possibly foodstuffs, may interact with immunity to parasites. Injury to adult *T. castaneum* induced the expression of genes for stress adaptation and insecticide resistance; thus, there may be crosstalk between immune and stress responses (Altincicek et al, 2008).

Resistance to pesticides and pathogens is thought to result in fitness costs, based on evolutionary models (Coustau et al, 2000). However, the insecticide-resistant strain of *T. castaneum* used in our studies had no or little, fitness costs considering responses to the parasite. Potential mechanisms of resistance to pathogens have been suggested (Coustau et al, 2000), but it is unclear how any of these could be related to the known malathion-resistance mechanisms (Coustau et al, 2000). The changes in ROS (including H_2_O_2_) could point to a common feature of both stressor types. Unfortunately, measurement of catalase in our *T. castaneum* strains exposed to *P. whitei* was not possible due to the low survival of the susceptible strain.

This is the first report on immunity in an insecticide resistant *Tribolium* sp. and to connect this to catalase activity. In this case, the strain has an uncommon resistance mechanism which involves a specific detoxication pathway. Our results suggest that pure oxygen and malathion exposure provoked oxidative stress and modulated catalase activity in the highly malathion-specific strain of *T. castaneum*, and an induction of a major antioxidant enzyme could have helped the strain to withstand both insecticidal and oxidative stresses. The results would seem to suggest that adaptation to toxic stress such as exposure to insecticides implies some changes other than the usual mechanisms (reduced uptake, modified target, enhanced metabolism) and these can induce major antioxidant enzymes that could have helped the strain to withstand both insecticidal and oxidative stresses.

