## supplementary materials for "Malathion-resistant *Tribolium castaneum* has enhanced response to oxidative stress, immunity, and fitness"

Supplementary Materials A

Insecticide toxicology of the strains used

Topical application: Acetone solution of each insecticide (0.5 µl/beetle) applied to ventral abdomen with a programmable Arnold microapplicator (Burkard) and a microsyringe (0.1ml) with a canula (G 36x3"). Acetone only was used for the control. Four replicates of ten insects each were used for each treatment, at room temperature and kept in an incubator for 24 hours at a temperature of 29±1°C and 65±5% relative humidity and then assessed. Treatments with synergists used the same procedure. The mortality data obtained were analysed by the probit method using a computer programme POLO PC (Leora Software, 1987) to obtain LD50, $\chi^2$ values, probits and heterogeneity factors.

Table 1 Toxicity of various insecticides to the strains of *T. castaneum* Herbst

| Strain | Insecticide | LD50 (95%CL) (µg/beetle) | Slope±SE | RF |
|---|---|---|---|---|
| Ph-1 | malathion | 6.4 (5-8) | 2.76±0.32 | 237 |
| | malaoxon | 1.7 (0.7-0.26) | 1.98±0.27 | 130 |
| | fenitrothion | 0.010 (0.006-0.015) | 3.12±0.44 | 2 |
| | permethrin | 0.047 (0.038-0.056) | 4.79±0.59 | 1.09 |
| | fenvalerate | 0.032 (0.029-0.035) | 5.4±0.69 | 1.03 |
| | methomyl | 0.023 (0.018-0.029) | 2.07±0.26 | 4 |
| | lindane | 0.98 (0.78-1.24) | 2.00±0.26 | 10 |
| | DEF (only) | 20.6 (19-22) | 7.54±1.02 | 2.3 |
| FSS-II | malathion | 0.027 (0.024-0.03) | 3.92±0.51 | |
| | malaoxon | 0.013 (0.08-0.023) | 2.709±0.31 | |
| | fenitrothion | 0.005 (0.005-0.006)) | 4.65±0.57 | |
| | permethrin | 0.043 (0.038-0.05) | 5.14±0.61 | |
| | fenvalerate | 0.031 (0.028-0.034) | 5.99±0.73 | |
| | methomyl | 0.006 (0.004-0.007) | 1.89±0.25 | |
| | lindane | 0.097 (0.073-0.125) | 1.76±0.25 | |
| | DEF (only) | 8.8 (8-9) | 5.19±0.67 | |

RF: Resistance factor=LD50 of resistant strain/LD50 of susceptible strain

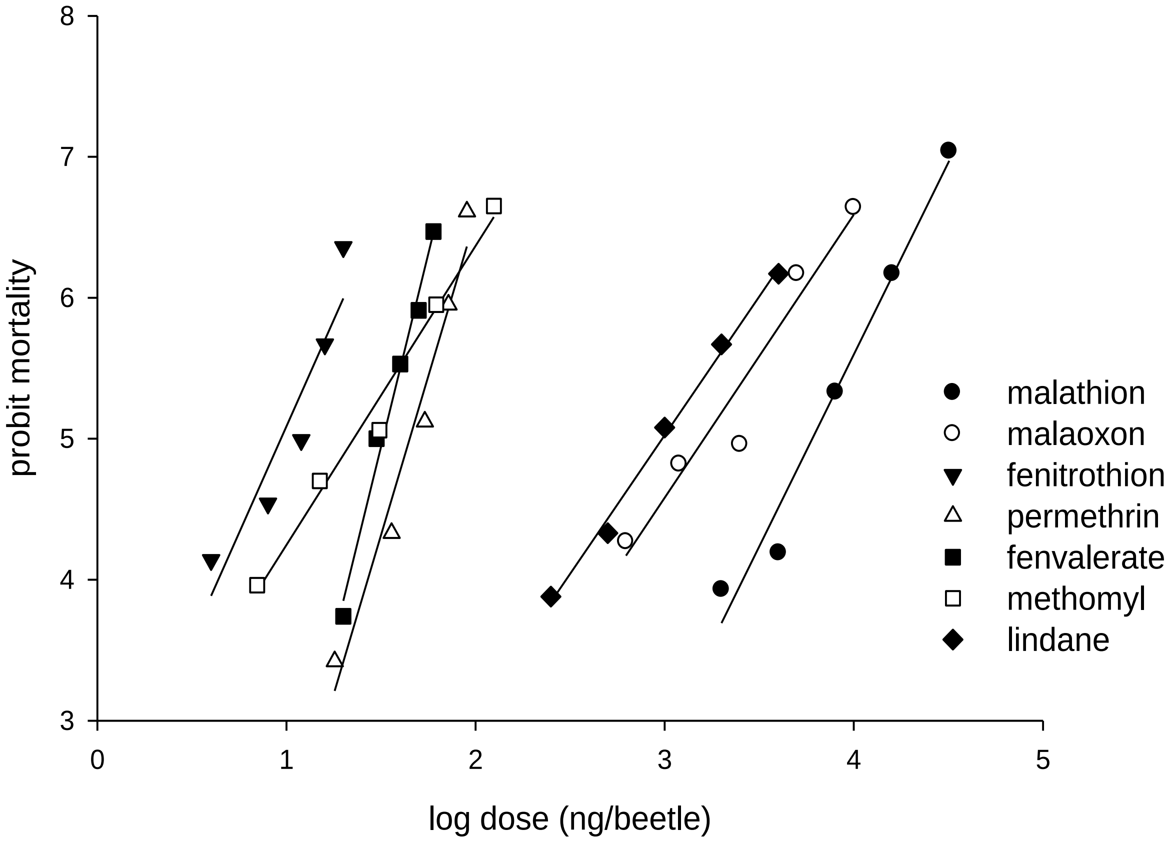


Fig. 1 Dose mortality response of Ph-1 strain to insecticides

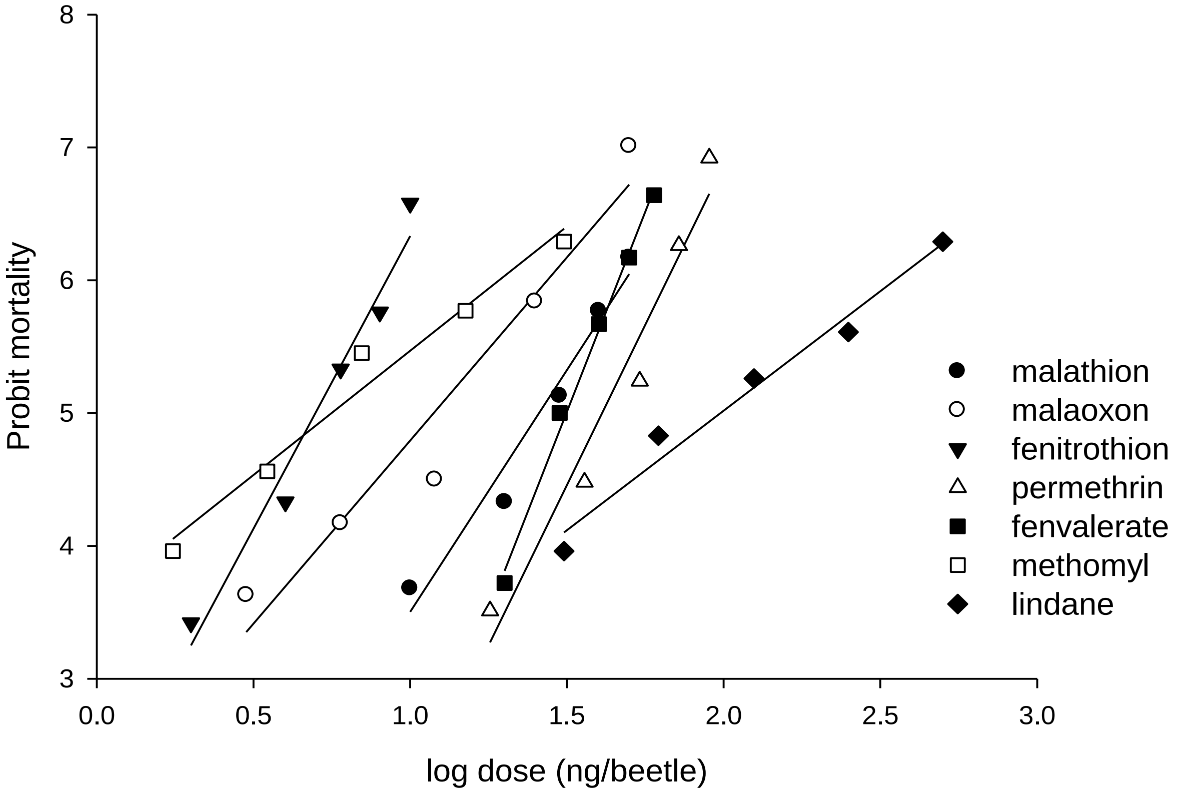


Fig. 2 Dose mortality response of FSS-II strain to insecticides

Table 2. Toxicities of insecticide + synergist (1:5) combinations to the strains of *T. castaneum*

| Strain | Synergist | LD50[a] µg/beetle | Slope ±SE | SF | $RF_1$ | $RF_2$ | $RF_2$/ $RF_0$ |
|---|---|---|---|---|---|---|---|
| PH-1 | malathion +DEF | 0.058 (0.04-0.07) | 4.06 ±0.51 | 110 | 2.1 | 2.6 | 0.01 |
| PH-1 | malathion+TPP | 0.083 (0.05-0.15) | 1.93 ±0.26 | 77 | 3.1 | 3.2 | 0.01 |
| PH-1 | malathion+PBO | 10 (7.6-13.4) | 1.48 ±0.19 | 0.64 | 370 | 385 | 1.6 |
| PH-1 | malathion+DEM | 14 (10-19) | 1.36 ±0.18 | 0.46 | 519 | 298 | 1.3 |
| FSS-II | malathion+DEF | 0.022 (0.02-0.026) | 3.27±0.44 | 1.2 | | | |
| FSS-II | malathion+TPP | 0.026 (0.022-0.029) | 3.35±0.46 | 1 | | | |
| FSS-II | malathion+PBO | 0.026 (0.021-0.032) | 3.84±0.49 | 1 | | | |
| FSS-II | malathion+DEM | 0.047 (0.04-0.05) | 3.99±0.49 | 0.6 | | | |
| PH-5 | malathion+TPP | 0.16 (0.12-0.23) | 2.45 ±0.26 | 843 | 3.3 | 3.6 | 0.001 |
| FSS-II (CSL) | malathion+TPP | 0.044 (0.03-0.05) | 3.88 ±0.48 | 1.1 | | | |

| | | | | | | | |
|---|---|---|---|---|---|---|---|
| PH-1 | malaoxon+DEF | 0.02 (0.015-0.024) | 2.18±0.27 | 85 | 1.5 | 4 | 0.03 |
| PH-1 | malaoxon+TPP | 0.026 (0.02-0.03) | 2.58±0.30 | 65 | 2 | 2.6 | 0.02 |
| FSS-II | malaoxon+DEF | 0.005 (0.004-0.007) | 1.83±0.25 | 2.6 | | | |
| FSS-II | malaoxon+TPP | 0.01 (0.007-0.013) | 2.19±0.27 | 1.3 | | | |
| PH-1 | lindane+PBO | 0.226 (0.14-0.42) | 1.66±0.20 | 4.3 | 2.3 | 2.4 | 0.24 |
| PH-1 | lindane+DEM | 0.892 (0.74-1.07) | 2.19±0.27 | 1.1 | 9.2 | 9.1 | 0.9 |
| FSS-II | indane+PBO | 0.094 (0.084-0.11) | 3.64±0.47 | 1 | | | |
| FSS-II | lilndane+DEM | 0.098 (0.08-0.12) | 1.96±0.26 | 1 | | | |

DEF: tributyl phosphorotrithioate, TPP: triphenyl phosphate, PBO: piperonyl butoxide, DEM: diethyl maleate. SF: LD50 without synergist divided by LD with synergist, $RF_1$: LD50 R strain with synergist / LD50 S strain without synergist, $RF_2$: LD50 R strain with synergist / LD50 S strain with synergist. [a] 95% confidence intervals.

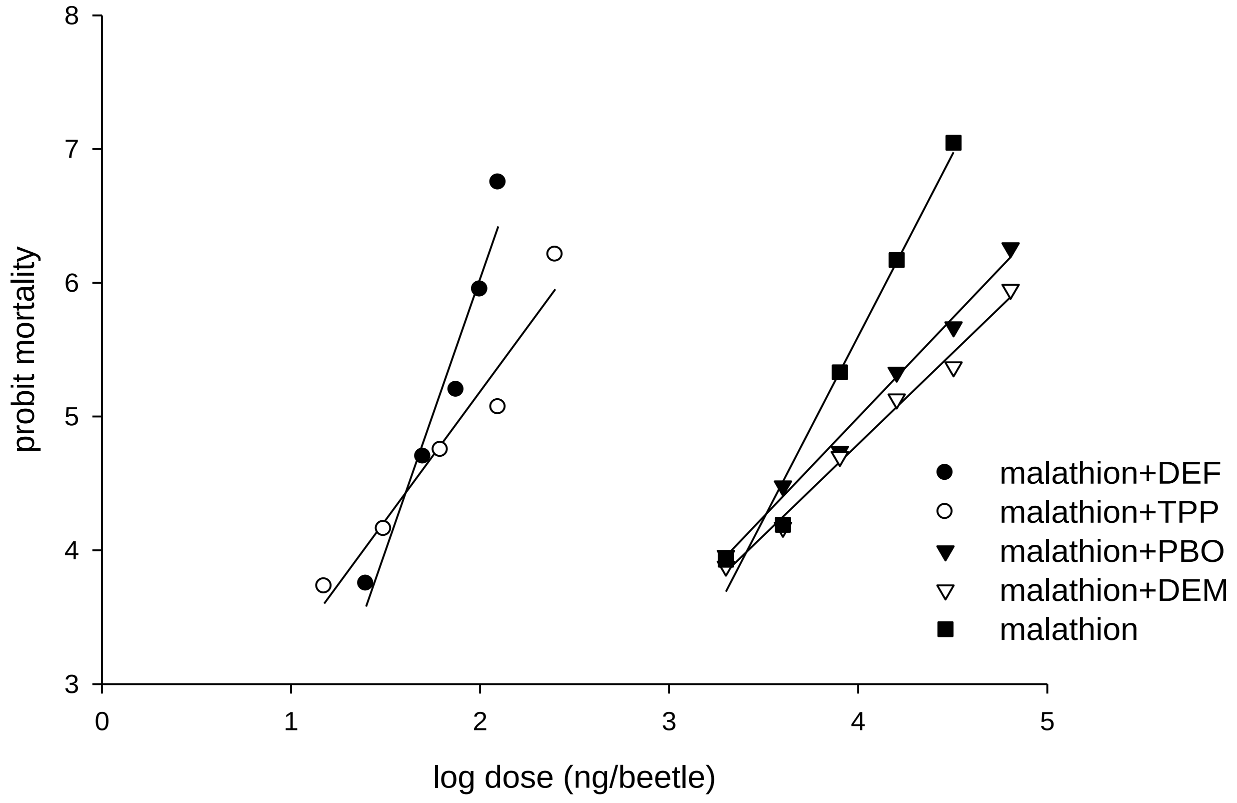


Fig. 3 Dose mortality response of PH-1 strain to malathion and malathion+synergists

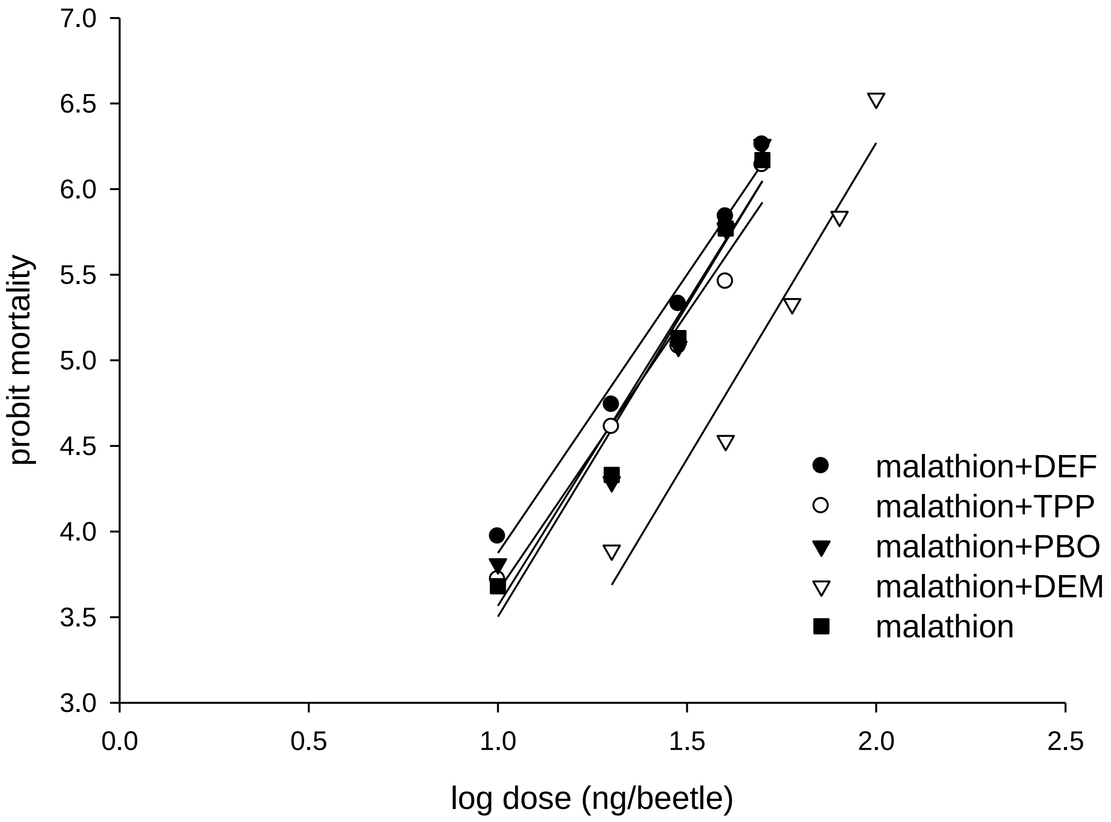


Fig. 4 Dose mortality response of FSS-II strain to malathion and malathion+synergists

Supplementary Materials B

Malathion metabolism in the two strains used

#### *in vivo* malathion metabolism in *T. castaneum*

Twenty-five beetles of each strain were first dosed with 12 mM triphenyl phosphate TPP solution in acetone using a topical application method 0.5 µl/beetle with acetone solvent. Another control set was topically treated with acetone alone. Then the same beetles were treated with ($^{14}$C-methoxy)-malathion, 5 µCi/ml in acetone using the same topical application method. The treated beetles were kept in the incubator at 30$^{o}$ C for 20 minutes. After the incubation period the insects were removed and washed twice with 1 ml acetone to remove any residual malathion. The group of beetles were then homogenized in 0.5 ml of 0.05M phosphate buffer (pH 7.6). The tubes were put in ice and 500 µl of n-hexane was added. The tubes were vortexed, shaken for 5 min with a shaker and then spun in a microcentrifuge for 1 min at 100000 rpm. The upper hexane layer was transferred directly into a scintillation vial containing 5 ml of scintillation cocktail. This extraction was repeated twice more and the extracted hexane being added to the same scintillation vial. Each homogenate was then acidified (to below pH 2) by the addition 50 µl of concentrated HCl. Subsequently three further extractions with 500 µl of diethyl ether were transferred directly to a scintillation vial containing 5 ml of scintillation cocktail. All scintillation vials were vortexed and the level of radioactivity was measured in a Packard liquid scintillation counter for 10 min. All measurements were made against a blank containing the cocktail only. Three replications were used for the assay and protein concentration in all homogenates was determined by the method of Bradford (1976) using BSA as standard protein.

**Statistical analysis**

The data was analysed by statistical programme GraphPad Prism 4, using two-way ANOVA with Bonferroni posttest.

**Results of *in vivo* malathion metabolism**

The resistant beetles, without TPP treatment, contained a lower mean level of malathion, after 20 min incubation but the difference was non-significant ($t(24) = 2.08$, $P>0.05$) as compared to the susceptible strain (Fig 1). The results also indicate that the total of malathion recovered in PH-5 strain was greater than of FSS-II but not significantly different.

The amount of metabolite (not specified) in the resistant strain Ph-5 was highly significantly greater than in the susceptible ($t(24) = 4.16$, $P<0.001$). However, in the presence of the inhibitor TPP metabolism was depressed in the resistant strain and was not significantly different to that in the susceptible FSS-II (Fig 1).

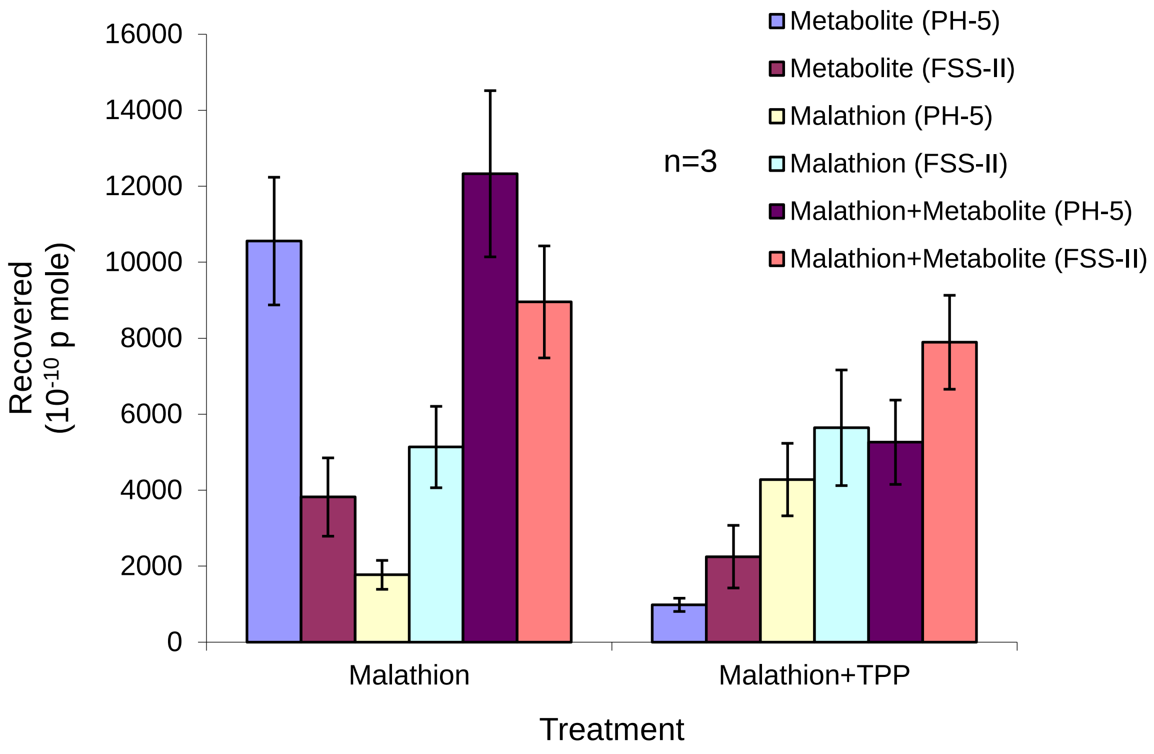


Fig. 1. *in vivo* malathion metabolism in the strains of *T. castaneum* (resistant Ph-5, susceptible FSS-II) n=number of replications, where, 25 beetles for each replication were used.

### *in vitro* malathion metabolism

#### Methods

With slight modifications, the method, followed by Spencer et. al (1998), was used for the quantitative analysis of *in vitro* malathion metabolism. According to that, fifty milligrams of resistant and susceptible adult insects were each homogenized in 2 ml ice cold 0.05 M phosphate buffer (pH 7.6), centrifuged at 11000 rpm and 4$^{o}$C in temperature controlled Eppendforf centrifuge for 10 min. The supernatant was transferred to a fresh tube and made up to 5 ml with the homogenizing buffer. Eight 500 µl aliquots were transferred to fresh tubes, four of which were placed in boiling water for 5 min and placed back in ice to cool. These four tubes will be used as control. 1.25 µl of ($^{14}$C-methoxy)-malathion (37.2 mM, 1.25 µCi ml$^{-1}$) was added to each tube and the tubes were kept in ice during the treatment. Then the treated tubes were incubated at 30$^{o}$ C for 15 min. Both homogenates were assayed for protein concentration by the method of Bradford (1976). The complete assay was carried out three times. After incubation, the tubes were again put in ice and 500 µl of n-hexane was added in each tube. The tubes were vortexed, shaken for 5 min with the help of a shaker and then spun in a microcentrifuge for 1 min at 100000 rpm. The upper hexane layer was transferred directly into a scintillation vial containing 5 ml of scintillation cocktail. This extraction was repeated twice more and the extracted hexane being added to the same scintillation vial. Each homogenate was then acidified (to pH <2) by the addition 50 µl of concentrated HCl. Subsequently three further extractions with 500 µl of diethyl ether were transferred directly to scintillation vial containing 5 ml of scintillation cocktail. To determine the residual activity 100 µl of each homogenate were added to a scintillation tube containing 5 ml of the cocktail. All scintillation vials were vortexed and the level of radioactivity was measured in a Packard liquid scintillation counter for 10 min. All measurements were made against a blank containing the cocktail only.

**Results**

At pH 7 for 15 min incubation the rate of metabolism of $^{14}$C-malathion was 0.257 (Ph-1) and 0.0091 (FSS-II) nmoles/min/mg protein ($t$ (16) = 4.41, $P<0.001$). Activity was completely inhibited by 2.2 nM inhibitor TPP. Rate of hydrolysis in the resistant strain was up to 28.17 times that in the susceptible strain.

Spencer, A. G., N. R. Price, and A. Callaghan. 1998. Malathion-specific resistance in a strain of the rust red grain beetle *Cryptolestes ferrugineus* (Coleoptera : Cucujidae). Bull Entomol Res 88: 199-206. https://doi.org/10.1017/S0007485300025761
